# Nucleolus association of chromosomal domains is largely maintained in cellular senescence despite massive nuclear reorganisation

**DOI:** 10.1101/054908

**Authors:** Stefan Dillinger, Tobias Straub, Attila Németh

## Abstract

Mammalian chromosomes are organized in structural and functional domains of 0.1-10 Mb, which are characterized by high self-association frequencies in the nuclear space and different contact probabilities with nuclear sub-compartments. They exhibit distinct chromatin modification patterns, gene expression levels and replication timing. Recently, nucleolus-associated chromosomal domains (NADs) have been discovered, yet their precise genomic organization and dynamics are still largely unknown. Here, we use nucleolus genomics and single-cell experiments to address these questions in human embryonic fibroblasts during replicative senescence. Genome-wide mapping reveals 1,646 NADs in proliferating cells, which cover about 38% of the annotated human genome. They are mainly heterochromatic and correlate with late replicating loci. Using Hi-C data analysis, we show that interactions of NADs dominate interphase chromosome contacts in the 10-50 Mb distance range. Interestingly, only minute changes in nucleolar association are observed upon senescence. These spatial rearrangements in subdomains smaller than 100 kb are accompanied with local transcriptional changes. In contrast, large centromeric and pericentromeric satellite repeat clusters extensively dissociate from nucleoli in senescent cells. We use gene set enrichment analysis (GSEA) to map the epigenetic regulatory network that governs these changes. The GSEA results together with cellular chromatin analyses suggest that histone H3K9 trimethylation is involved in regulating the nucleolus association of chromatin. Collectively, this study identifies connections between the nucleolus, 3D genome structure, and cellular aging at the level of interphase chromosome organization.

## Introduction

The spatiotemporal regulation of genomes correlates with transcription, replication, recombination and repair. In the post-genomic era a new model of human genome organization emerged, which is largely based on high-throughput genomics analyses. A key concept of this model is that chromosomal domains, megabase-ranged functional units of the chromosomes, represent an essential operational level of genome regulation (reviewed in ^1-8^). However, several questions remained open about the domain organization of human chromosomes. These include the dynamics and functional consequences of chromosomal domain interactions with nuclear sub-compartments in different cell types and under various physiological conditions. The nucleolus is a paramount example for the functional organization of the genome in space and time. Active nucleolar organizer regions (NORs) from different chromosomes build in all cell types a microscopically visible, dynamic nuclear compartment after mitosis, which disassembles again during the next cell division. Consequently, nucleolus-associated chromatin ^9^ also undergoes cyclic changes in proliferating cells. While the involvement of the nucleolus in ribosome biogenesis, furthermore in facilitating cell cycle progression, stress sensing and RNP function is well characterized ^10,11^, we just begin to uncover the molecular characteristics of the nucleolus-associated chromatin and to understand the role of the nucleolus in genome organization and function ^12-14^. Nucleolus-associated chromosomal domains (NADs) represent the mappable genomic fraction of the nucleolus-associated chromatin. NADs were first identified in HeLa cervical carcinoma and HT1080 fibrosarcoma cells using the combination of high-throughput genomics and immuno-FISH analyses ^15,16^ These studies provided us with a snapshot of global genome organization in and around the nucleolus and revealed that NADs represent mainly, but not exclusively a specific heterochromatin compartment. In addition, the mechanistic role of several *cis-* and trans-acting factors in chromosomal domain - nucleolus interactions has also been initially addressed ^17-20^. However, comprehensive high-resolution maps of NADs and the characterization of the nucleolus-associated genome of normal diploid cells have not been determined. Moreover, the involvement of NADs in the formation of interphase chromosome structure and its dynamics during diverse cellular processes such as cellular aging, differentiation or cell cycle, remained largely unknown.

Cellular senescence is a stable arrest of cell proliferation, whose common form, replicative senescence, is induced by telomere attrition and chromosomal instability. In human and other multicellular organisms, cellular senescence plays a pivotal role in several physiological processes, namely tumour suppression, tissue repair, embryonic development and organismal aging ^21,22^. Notably, senescence-related, genome-wide reprogramming of gene expression is accompanied by massive structural reorganization of chromatin ^23-28^ Changes in the spatial organization of chromosomes include the formation of senescence-associated distension of satellites (SADS), an early and consistent marker of cellular senescence ^29^, which can be followed by the development of senescence-associated heterochromatin foci (SAHF) under certain conditions ^30,31^. Remarkably, SAHF-like chromosome condensation can be induced also in a senescence-independent manner ^32^ The nucleolar hallmarks of senescent cells include increased size and fusion of nucleoli in mammals, furthermore instability of ribosomal DNA (rDNA) in yeast (reviewed in ^33,34^). Notably, the instability of the rDNA cluster is considered as a key control element of genome maintenance and inducer of cellular senescence in yeast ^35,36^ While recent works highlighted the involvement of various histone modifications, lamins and DNA methylation in shaping the epigenetic landscape during cellular senescence in mammals (reviewed in ^37^"^40^), the molecular details of senescence-related nucleolar remodelling are still elusive.

Here we present high-resolution NAD maps of young and replicative senescent IMR90 human embryonic fibroblasts. Our findings identify a central role for the nucleolus in the nuclear arrangement of specific transcriptionally silent chromosomal domains in proliferating and senescent cells, and delineate molecular mechanisms involved in the regulation of nucleolus-associated chromatin and spatial genome organization.

## Results

### Mapping and genomics of NADs in primary human cells

In two independent experiments, nucleoli of each young and proliferating IMR90 fibroblasts were isolated by biophysical disruption of cells and subsequent fractionation by differential centrifugation. To assess the quality of the purification, the enrichment of the nucleolar transcription factor UBTF and ribosomal DNA (rDNA) were monitored in immunoblot and quantitative PCR experiments, respectively, and the nucleolar fraction was examined also by microscopy (Supplementary Fig. S1). The DNA of isolated nucleoli was extracted and subjected to comparative genomic hybridization experiments on whole genome tiling microarrays to identify the non-repetitive DNA content of the nucleolus-associated genome and to address its chromosomal organization. The bimodal nature of hybridization signals provided the basis for genome-wide mapping of NADs with the use of a two-state hidden Markov model (HMM) analysis. By using this approach 1,646 autosomal NADs of young, proliferating IMR90 cells were discovered (Fig. 1a, Supplementary Fig. S2, and Supplementary Table S1). We refer here to this comprehensive list of IMR90 NADs simply as 'NADs', and to the previously identified HeLa NADs ^15^, which were mapped using an 85%-threshold-based method, as 'top NADs'. NADs cover 1.2 billion bp, approximately 38% of the annotated human genome, and their median sequence length (361 kb) resides in the typical size range of the higher order structural and functional chromosomal domains of mammalian genomes (Fig. 1b).

**Figure 1.**
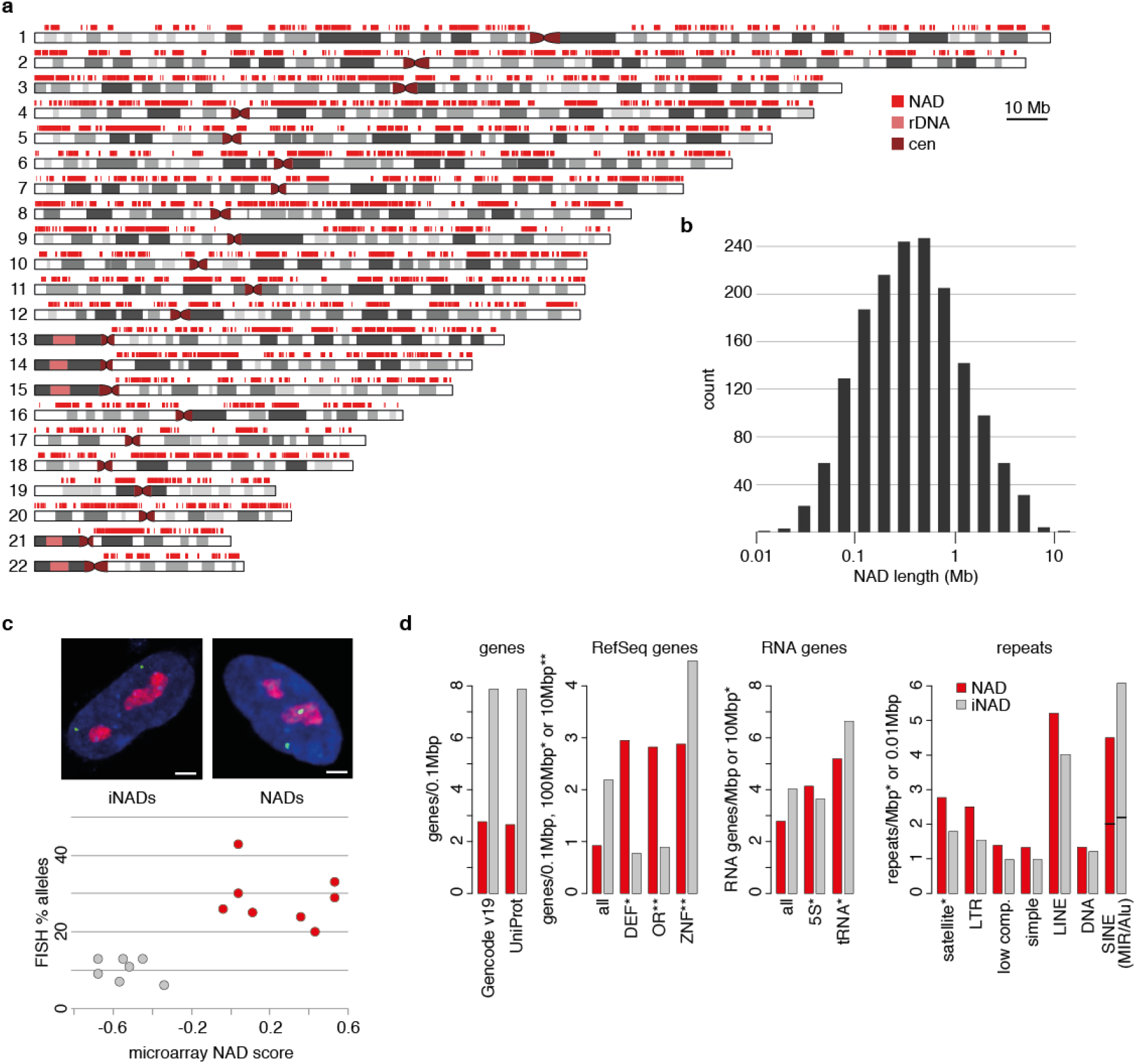
**Map and genome features of nucleolus-associated chromosomal domains (NADs) in IMR90 primary human embryonic fibroblast cells.** (**a**) Distribution of NADs along human autosomes. NADs are indicated by red rectangles over the ideograms of the chromosomes. Note that the p-arms of the five acrocentric chromosomes (13, 14, 15, 21 and 22), centromeres and some pericentromeric regions were not analysed because they are not present in the current human genome assembly. (**b**) Histogram of NAD sizes. Median=361kb, a total of 1,646 NADs were identified. (**c**) 3D immuno-FISH analysis of NAD and inter-NAD regions (iNADs) in IMR90 cells. The hybridization signal (percentage of nucleolus-associated alleles) is plotted against the according microarray signal (average log2-fold difference of the nucleolar signal over the background). Red and grey circles indicate genomic regions that reside in NADs and iNADs, respectively (see Table S2 for further details). Representative single light optical sections of IMR90 nuclei are shown on the top. BAC hybridization signals are shown in green, nucleolar staining in red and DAPI counterstain in blue (scale bars: 1.6 μm). (**d**) Bar graphs show Gencode v19, UniProt, RefSeq gene (ZNF, OR and DEF indicate zinc finger, olfactory receptor and defensin gene families, respectively), non-coding RNA gene (‘RNA genes’) and repeat frequencies in NADs (red) and iNADs (grey) based on UCSC Table Browser data.

In order to validate the nucleolus association of selected chromosomal regions, 3D immuno-FISH experiments were performed. Nucleoli were labelled by immunofluorescence staining of nucleophosmin (NPM1) and different genomic regions by fluorescence hybridization. The frequency of nucleolus interaction was determined for several genomic regions and plotted against the NAD score value, which was calculated from the microarray data by averaging the log-enrichment values of two replicate experiments. Additionally, our previously collected 3D immuno-FISH data ^15^ was also integrated in the analysis (Fig. 1c and Supplementary Table S2). The results of population-based and single-cell analyses correlate well, as chromosomal regions with high NAD scores showed typically more frequent association with nucleoli in 3D immuno-FISH than regions with low NAD scores.

Next, the incidence of various sequence features was addressed in IMR90 NADs and inter-NAD regions (iNADs) of the annotated human genome (Fig. 1d). The results revealed that defensin (DEF) and olfactory receptor (OR) genes are enriched in otherwise gene-poor NADs, and they are depleted in gene-rich iNADs. Although zinc finger (ZNF) genes are depleted in NADs, the extent of their depletion is less than the overall depletion of protein-coding genes in NADs. Non-coding RNA genes are also less prevalent in NADs than in iNADs, however 5S RNA genes are slightly enriched in NADs. Repetitive DNA analyses showed that satellite repeats and LTR elements are enriched in NADs, and to a far lesser extent also low complexity repeats, simple repeats, LINEs and DNA repeats. In contrast, SINEs are depleted in NADs due to the lower incidence of Alu repeats in these genomic regions. Taken together, the prevalence of specific sequence features in NADs is similar to that of top NADs, but the enrichment of individual features is less pronounced.

### Comparative functional epigenomics of NADs and iNADs reveals specific heterochromatic features of NADs and identifies X escapers as iNADs

In order to uncover the chromatin features of NADs at the genome level, the 15-state chromatin HMM (ChromHMM) map of IMR90 cells ^41^ was quantitatively analysed. The size of NAD and iNAD genomic regions occupied by the various chromatin states was identified and illustrated on a bar graph. Next, the values of each chromatin state were calculated for the entire population of NADs and iNADs, and their log2 ratios were plotted (Fig. 2a). The results provide compelling evidence that NADs are depleted in active chromatin features and they can be mainly characterized by the 'heterochromatin' and 'quiescent/low' ChromHMM states. The main features of these chromatin states are high levels of DNA methylation, low DNaseI accessibility, and low incidence of genes, which are mainly repressed. Further investigations of the overall DNaseI accessibility and transcriptional activity in NADs and iNADs revealed large differences with clearly lower values in NADs (Fig. 2b and 2c), supporting the results of the ChromHMM analysis.

**Figure 2.**
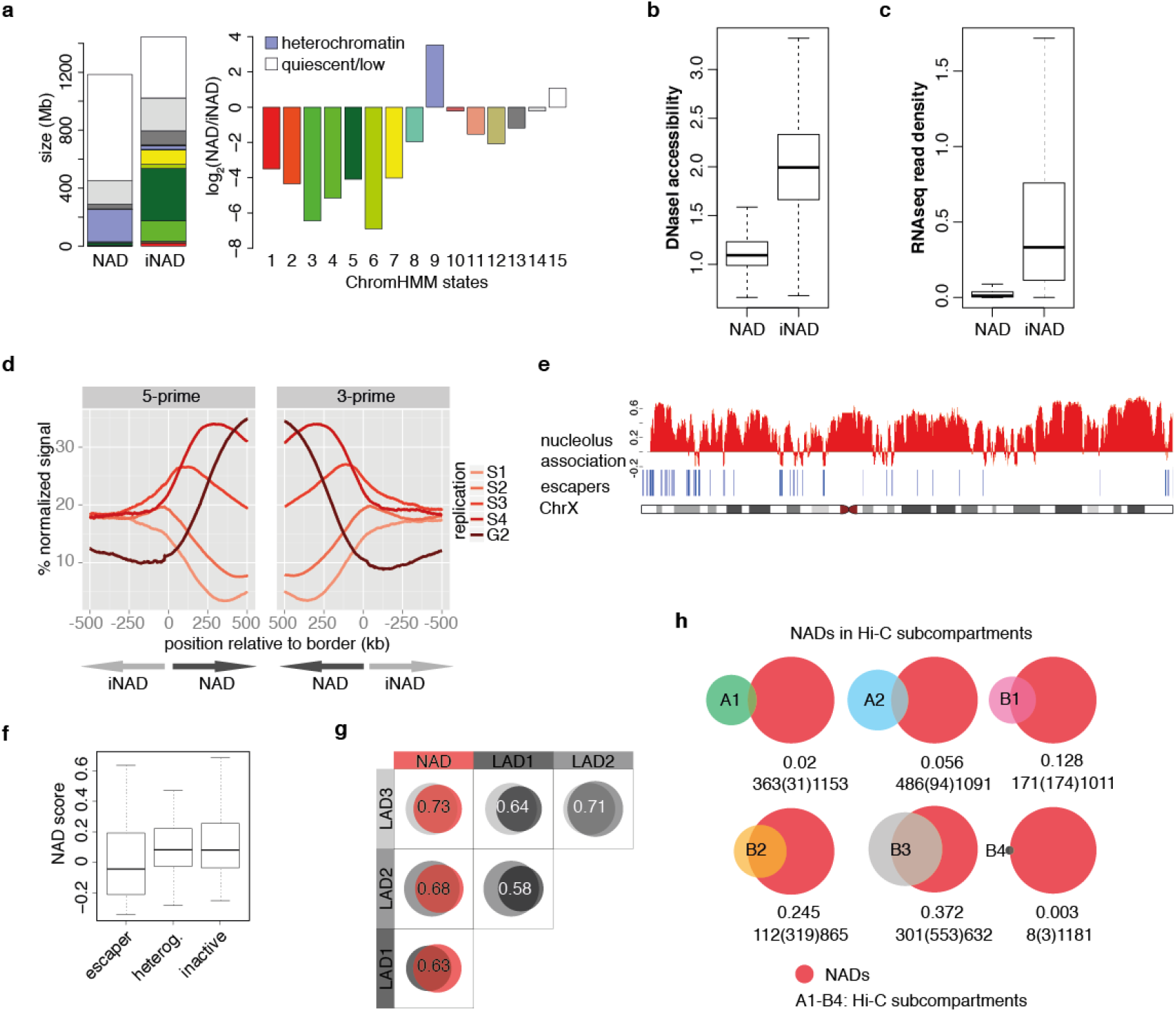
**Comparative functional epigenomics of NADs and iNADs reveals specific heterochromatic features of NADs and identifies X escapers as iNADs.** (**a**) Distribution of different chromatin states in NADs and iNADs. Bar graphs show total and relative amounts of ChromHMM states in NADs and iNADs on the left and right, respectively. The chromatin states and their colour code correspond to the Primary Core Marks segmentation 15-state ChromHMM model of the Roadmap Epigenomics Project^41^. Chromatin states specifically enriched in NADs are indicated. Red: Active Transcriptional Start Site (TSS), OrangeRed: Flanking Active TSS, LimeGreen: Transcription at gene ends, Green: Strong transcription, DarkGreen: Weak transcription, GreenYellow: Genic enhancers (Enh), Yellow: Enh, MediumAquamarine: ZNF genes & repeats, PaleTurquoise: Heterochromatin, IndianRed: Bivalent/Poised TSS, DarkSalmon: Flanking Bivalent TSS/Enh, DarkKhaki: Bivalent Enh, Silver: Repressed PolyComb, Gainsboro: Weak Repressed PolyComb, White: Quiescent/Low. (**b**) Boxplot of DNaseI accessibility in NADs (n=1646) and iNADs (n=1669). The average accessibility per base and segment were calculated from GSM468792. (**c**) Boxplot of RNA-seq read densities. For each NAD (n=1646) and iNAD (n=1669) the average number of reads per base in GSM438363 were calculated. (**d**) Replication timing profiles around NAD borders. Average percentage-normalized signals of different replication domains within a distance of 500 kb from aligned 5' and 3' NAD borders. Repli-Seq signals from five different time points of the cell cycle (S1-S4 and G2) were taken from Pope *et* al.,^4^ and averaged over 1 kb windows. Only NADs with a width >500 kb were considered (n=652). (**e**) Microarray signals of nucleolus-association (log2-fold difference of the nucleolar signal over the background, combined from two biological replicate experiments) and X escaper gene positions are shown on the top of the X chromosome ideogram. (**f**) Boxplot of NAD scores in 'escaper', 'heterogeneous' and 'inactive' genes (classification based on Carrel and Willard^43^). (**g**) Venn diagrams and Jaccard coefficients show the extent of overlap between NADs and LADs. LAD1: LADs of Tig3 cells^45^, LAD2 and LAD3: LADs of IMR90 cells^24,25^. (**h**) Venn diagrams of NADs and A1-A2, B1-B4 Hi-C subcompartments show heterochromatic features of NADs. Jaccard coefficients and the sizes of overlapping and non-overlapping regions (in Mb) are shown below the diagrams.

To identify the replication timing profile around NAD borders, NADs were aligned at their 5' and 3' ends (p to q chromosome orientation), and normalized Repli-seq signals were averaged in 500 kb windows spanning the borders. Replication signals were measured recently at five different time points of the S/G2 phases of the cell cycle ^42^, and their distributions appear as five waves around NAD borders (Fig. 2d). The waves show a regular spatiotemporal pattern within NADs, in which early-to-late replication timing correlates with the distance to the border. Interestingly, only the depletion of the latest replicating fraction (G2) is notable outside of NADs in the 0 to −500kb distance range, whereas the signal intensities of the other four fractions (S1 to S4) get indistinguishable with increasing distance. Altogether, the result is in good agreement with the late-replicating nature of heterochromatin, the characteristic chromatin state of NADs.

As IMR90 cells possess one active and one inactive X chromosome having substantially different chromatin conformations and no Y chromosome, the sex chromosomes were not included in the aforementioned genome-scale NAD analyses. In an extended mapping of NADs a few regions of the Y chromosome emerged as nucleolus associated, due to cross-hybridization or false assignment on the microarrays. In contrast, almost the entire X chromosome appeared as NAD in this analysis. The heterochromatic nature of the inactive X chromosome and the frequent association of the entire chromosome with the perinucleolar heterochromatin might explain this result (Supplementary Fig. S3). Nevertheless, certain regions of the X chromosome were identified as iNADs. Since some of them clearly overlapped with the pseudo-autosomal regions that escape X inactivation (Fig. 2e), the correlation between escaper genes and iNADs of the X chromosome was further addressed. The 'escaper', 'heterogeneous' and 'inactive' states of genes were classified according to Carrel and Willard (^43^ and Supplementary Table S3) and NAD score values were computed for all three groups. A boxplot representation of the results demonstrates that the escaper group is less frequently associated with nucleoli than the two others (Fig. 2f). The visualization of microarray data along with escaper gene positions provides a more detailed picture. Besides the fact that nearly all iNAD regions contain escaper genes, the remaining escapers also show a strong coincidence with weaker nucleolus association within NADs (Fig. 2e). These observations strongly suggest that local spatial distension, for instance looping of active genes leads to their efficient separation from the more compact heterochromatin of the inactive X chromosome during nucleolus isolation.

As described above, several lines of experimental evidence point to a considerable enrichment of heterochromatin in NADs. The constitutive heterochromatin of cultured mammalian cells predominantly localizes to the nuclear periphery, pericentromeric bodies and the perinucleolar region, and these nuclear sub-compartments are to large extent functionally overlapping (recently reviewed in ^44^). Genomic analyses of the nuclear periphery identified the maps of lamina-associated domains (LADs) in proliferating Tig3 human fibroblasts by using the DamID method ^45^, and later also in IMR90 cells by a Lamin B1 ChIP-seq approach ^24,25^ To statistically evaluate the similarity of LADs and NADs of human embryonic fibroblasts, the three LAD datasets were compared to the NAD list of IMR90 cells and to each other in a quantitative manner. As illustrated by the large intersections of the Venn diagrams, the IMR90 NADs show a substantial overlap with the LADs of human diploid fibroblasts (Fig. 2g). However, about one-third of the total NAD- and LAD-covered genomic regions are non¬overlapping and chromosome-specific differences in the patterns are clearly recognizable (Supplementary Fig. S4).

Chromosome Conformation Capture (3C) provides a way to divide chromosomes into domains by measuring contact probabilities between chromosomal segments. Hi-C analyses with increasing resolution led to the genome-wide determination of A and B compartments ^46^, topological domains ^47^, and finally contact domains ^48^ of human chromosomes. However, this methodology does not deliver information about the nuclear position of chromosomal domains and their interaction with nuclear bodies. To integrate complementary information about the spatial organization of the human genome in IMR90 cells, a comparative genomic analysis was performed using the NAD data and the to date highest resolution Hi-C datasets of this cell line ^48^. Venn diagrams show the overlap between NADs and the six genomic subcompartments, which were determined by Hi-C-based segregation of the human genome (Fig. 2h). First, the results demonstrate that there is only little overlap between NADs and the euchromatic, gene dense A1-A2 subcompartments. Second, nearly half of the B1 subcompartment, which represents facultative heterochromatin, coincides with NADs. Third, approximately three-quarter of the B2 and two-third of the B3 subcompartments, primarily constitutive heterochromatin, correspond to NADs. Importantly, these are the largest Hi-C subcompartments. Of note the strongest overlap between B2 and the NADs, which was also detected with the top NADs ^48^, and also the increased B3 - NAD overlap. Finally, NADs coincide to a greater extent with B4 than with A subcompartments. The interpretation of this overlap is ambiguous due to the minute size of B4. B4 contains KRAB-ZNF gene clusters and displays a specific chromatin pattern, and it is the far smallest Hi-C subcompartment. Altogether, the results of the comparative analysis of Hi-C subcompartments and NADs are largely consistent with the results of the genomic analyses shown in Fig. 2a-d, and uncover that 74% of NADs reside in B2/B3-type constitutive heterochromatic chromosomal regions.

### NAD-NAD interactions dominate in the 10-50 Mb distance range over iNAD-iNAD and heterotypic NAD-iNAD interactions

In the nucleus of mammalian cells the chromosomes occupy distinct territories ^1,49,50^, and their ultrastructural organization is influenced by intra-chromosomal association of chromosomal domains. To compare the contribution of NADs and iNADs to intra-chromosomal interactions at different size scales, frequencies of NAD-NAD, iNAD-iNAD and mixed NAD-iNAD contacts were calculated for each chromosome from IMR90 Hi-C datasets and plotted against linear sequence distance (Fig. 3a and Supplementary Fig. S5). Intra- and interarm interactions were visualized separately to consider a possible influence of centromeres on intrachromosomal contact probabilities. Visual inspection of the plots revealed that NAD-NAD contacts dominate at the 10-50 Mb distance range over iNAD-iNAD and mixed contacts. Over larger distances iNAD-iNAD contacts became the most frequent, whereas NAD-iNAD contact probabilities are clearly the lowest ones at all distance ranges. Moreover, a strong increase of contact frequencies can be observed in the <10 Mb distance range, which corresponds well to the size range of topologically associating domains (TADs). TADs are considered as the basic units of chromosome folding and they can be defined by measuring genomic interaction frequency changes along the chromosomes: the interaction frequencies of two loci are high within TADs but sharply drop at the boundary between neighbour TADs ^51^. In order to visualize NAD-NAD and iNAD-iNAD contacts at >10Mb distances in a locus-specific manner, the Hi-C contact frequencies of individual chromosomes were displayed also on heat maps (Supplementary Fig. S6). To aid the identification of homotypic contacts, NADs or iNADs were masked in additional individual heat maps. Strikingly, NAD/iNAD and Hi-C-contact-based TAD segmentations of the chromosomes revealed highly similar patterns. In addition, the heat maps showed again that NAD-NAD contacts are enriched at 10-50 Mb and depleted at >50 Mb distances.

**Figure 3.**
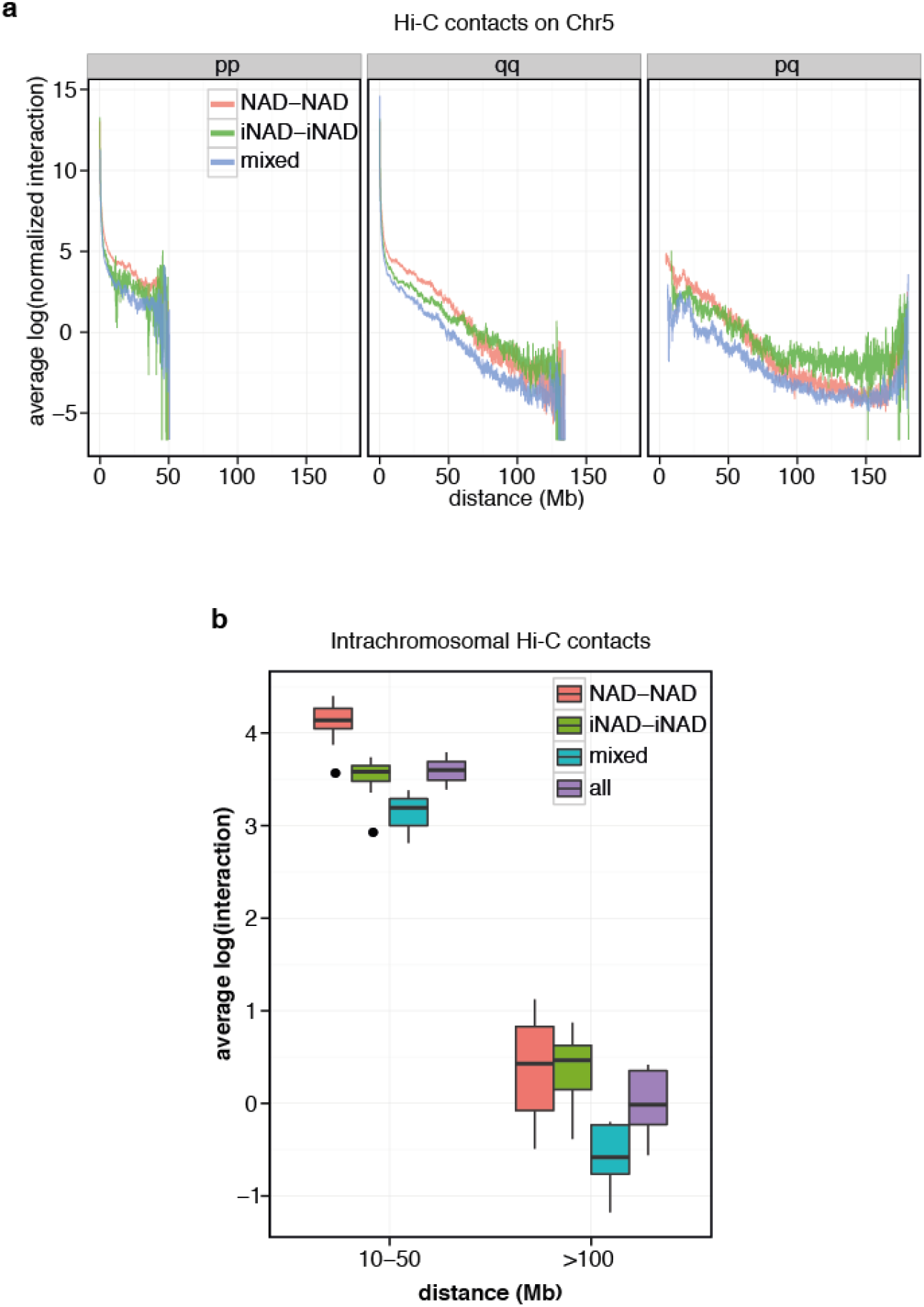
**NAD-NAD interactions dominate in the 10-50 Mb distance range over iNAD-iNAD and heterotypic NAD-iNAD interactions.** (**a**) Average intrachromosomal NAD-NAD, iNAD-iNAD and NAD-iNAD interactions at different distances on chromosome 5. (**b**) Boxplots of genome-wide NAD-NAD, iNAD-iNAD and NAD-iNAD interaction frequencies measured at 10-50Mb and >100Mb distances. Data points are the average interaction values of 10 kb windows matching a specific interaction class (NAD-NAD, iNAD-iNAD, mixed) and calculated for chromosomes 1 to 12 (n=12).

Next we calculated the average interaction intensity within each NAD-NAD, iNAD-iNAD, and NAD-iNAD contact to address the involvement of NADs in chromosomal contacts at different distance ranges. To exclude intra-NAD contacts from the analyses, contact frequencies at <10Mb distances were omitted. The contact frequencies of different contact classes were calculated at two different distance ranges, at 10-50Mb versus >100Mb and shown in a boxplot diagram (Fig. 3b). Due to the <100Mb size of human chromosomes 16 to 22 and the fact that the p arms of the acrocentric chromosomes 13 to 15 are not annotated, only chromosomes 1 to 12 were investigated here. The boxplots show that homotypic interactions of chromosomal domains are the prominent ones in the organization of chromosome territories at >10Mb distances. Importantly, the results also reveal that NAD-NAD contacts are the dominant ones at the 10-50Mb distance range. Next, the contribution of the different contact classes to the overall contact frequencies within and between chromosome arms was visualized (Supplementary Fig. S7). It is clearly recognizable that NAD-NAD interactions are the dominant ones within intraarm contacts over 10Mb distances, whereas NAD-NAD and iNAD-iNAD interactions show similar frequencies within interarm contacts. The prevalence of NAD-NAD interactions in the intraarm contacts compared to interarm contacts suggests that the centromere may reduce interarm contact probabilities, however the linear distance seems to play the major role in determining intrachromosomal NAD-NAD contact probabilities.

### Replicative senescence causes fusion and enlargement of nucleoli, but only local, transcription-dependent changes in NADs

Next, we addressed the questions how nucleolar morphology, NAD organization and global gene expression alter during cellular aging. IMR90 fibroblasts were grown under standard cell culture conditions and the aging of the cell population was monitored (Supplementary Fig. S8). In order to evaluate senescence-associated changes in nucleolar number and morphology, proliferating and senescent IMR90 cells were stained for NPM1 by immunofluorescence. Counting of NPM1 signals showed that proliferating cells exhibit typically 2-4 nucleoli and senescent cells a single, large nucleolus (Fig. 4a). In addition, nuclear and nucleolar volumes were measured to quantitatively assess the senescence-associated large-scale reorganization of the nucleolar subcompartment in the human nucleus. The results showed that the median volume of individual nucleoli in senescent IMR90 cells is about 6-fold larger than in young, proliferating cells. Moreover, the total nucleolar volume per cell also increases about 2.6-fold in senescence. This is not only due to the enlargement of the nucleus, as the relative median volume that is occupied by nucleoli also increases by more than 2-fold when cells enter senescence (Fig. 4b).

**Figure 4.**
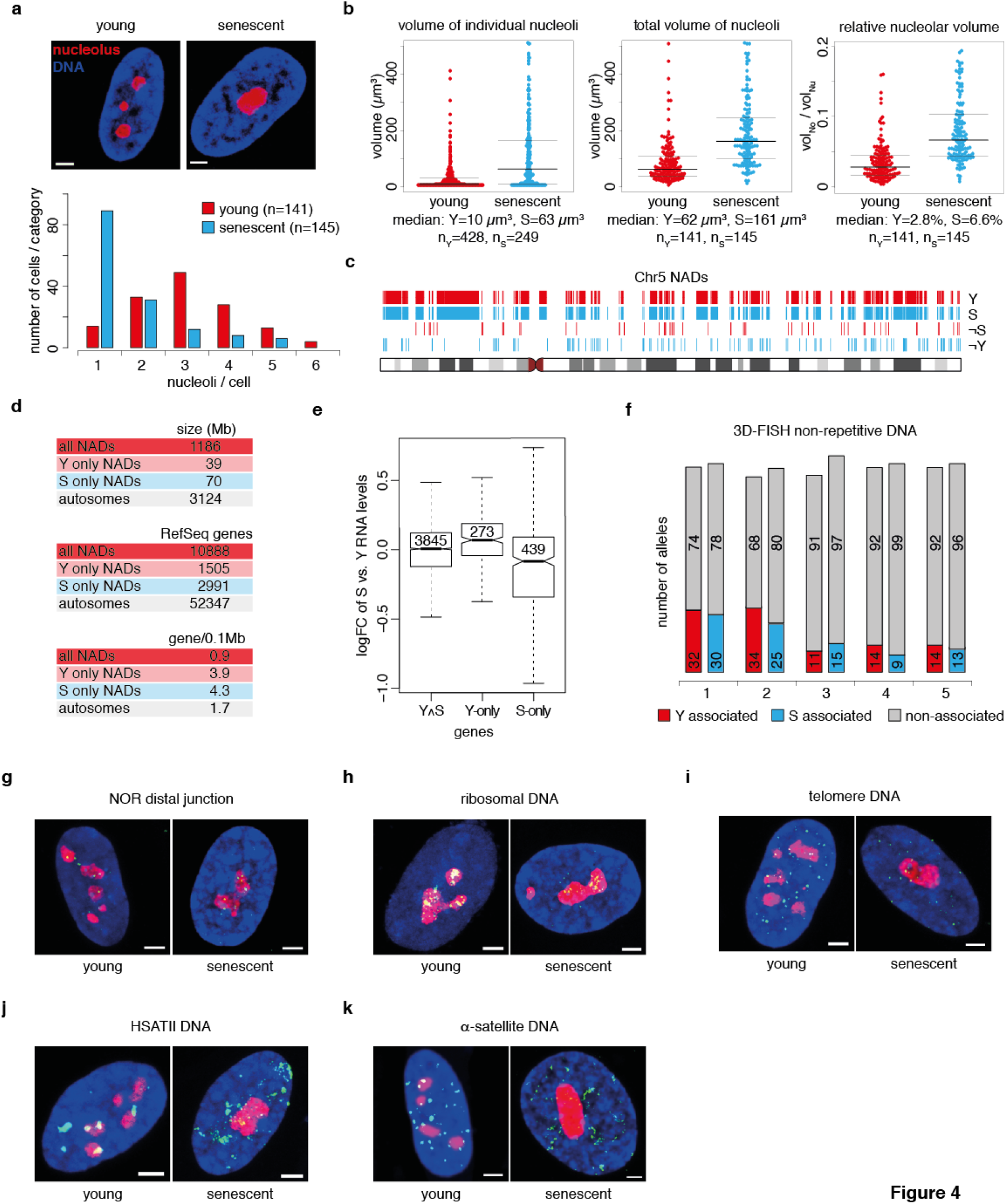
**Replicative senescence causes fusion and enlargement of nucleoli, but only local, transcription-dependent changes in NADs, furthermore it impairs nucleolus - satellite repeat cluster interactions.** (**a**) Bar graph of nucleolus number in young, proliferating ('Y') and senescent ('S') IMR90 cells. Proliferating cells have 3.0±1.2 and senescent cells 1.7±1.1 nucleoli per nucleus. Mid-sections of representative confocal microscopy images are shown on the top. Nucleolar staining is shown in red and DAPI counterstain in blue (scale bars: 1.6 μm). (**b**) Bee swarm plots show the volume of individual nucleoli, the total volume of nucleoli per nucleus and the nucleolar/nuclear volume ratio as measured by nucleolar immunostaining and DAPI staining of proliferating and senescent IMR90 cells. Center lines show the medians; upper and lower thin lines indicate the 25^th^ and 75^th^ percentiles. (**c**) Maps of NADs on chromosome 5 from young and senescent cells. Genomic regions associated with nucleoli only in young (-S) or senescent (-Y) cells are shown also as individual tracks. (**d**) Y-only and S-only NADs are enriched in protein-coding genes compared to all NADs and the genome. RefSeq gene data were obtained from the UCSC Table Browser. (**e**) Boxplots show positive correlation of senescence-related loss of nucleolus association and gene activation. Global gene expression changes (log2 fold change in senescent versus young cells) in constitutive (YAS), Y-only and S-only NAD genes are shown. The notches are defined as +/-1.58*IQR/sqrt(**n**) and represent the 95% confidence interval for each median. Group means are significantly different for all comparisons (p-value < 0.05, T ukey HSD). (**f**) Stacked columns show that the association frequency of five selected genomic regions is similar in young and senescent cells. Nucleolus-association data were collected from 50 cells for each category. BAC clones 1 to 5: RP11-44B13, RP11-173M10, RP11-828F4, RP11-125O21, RP11-81M8. (**g**) The distal junction region of NORs remains associated with nucleoli in senescent cells. Mid-sections of representative confocal microscopy images are shown (in g to k). Nucleolar staining is in red, DAPI counterstain in blue and FISH signals are in green on all images (scale bars: 1.6 μm). (**h**) Localization of the rDNA repeat clusters remains also nucleolar in senescent cells. (**i**) The signal intensity of telomeres is reduced in senescence, but their overall association with nucleoli does not show specific changes. (**j**) HSATII and (**k**) alpha-satellite repeat clusters display senescence-associated distension and reduced nucleolus association.

In order to discover genome-wide changes of nucleolus-associated chromatin at high resolution, NADs of senescent cells were mapped and compared to NADs of young, proliferating cells. Surprisingly, the two NAD maps were highly similar. Altogether 39 Mb sequence was specific for young NADs and 70 Mb for senescent NADs, which respectively represent 3.3% and 5.9% of the total NAD size (1.2 billion bp) determined in young IMR90 fibroblasts (Supplementary Table S4). Importantly, chromosomal fragments that can be determined as young-only or senescent-only NAD regions have a median size of less than 20 kb. Accordingly, most of the changes involved just parts of individual NADs (Fig. 4c and Supplementary Fig. S9). Systematic analysis of sequence features in young-only and senescent-only NAD regions revealed that they are particularly enriched in protein-coding genes (Fig. 4d and Supplementary Table S4). To identify whether the changes in nucleolus association correlate with transcriptional dynamics gene expression microarray experiments were performed. A targeted bioinformatics analysis revealed that the loss of nucleolus association correlates with an overall significant increase in transcript levels, while the gain of nucleolus association with a significant decrease of it (Fig. 4e). Additionally, hierarchical clustering analysis of gene expression microarrays was performed using the data from this and two other recent studies ^25,52^, in which the same experimental system was investigated. The result showed co-clustering of young and senescent samples providing an additional, robust quality control of the proliferating and senescent states (Supplementary Fig. S10).

### Nucleolus - satellite repeat cluster interactions are impaired in senescent cells, whereas rDNA, telomeres and single copy regions do not display remarkable changes in nucleolus association frequency

To uncover the role of cellular aging on the nuclear position of selected genomic loci 3D immuno-FISH analyses were performed. The comparison of the association frequencies of five chromosomal loci with nucleoli showed only little differences between young and senescent cells (Fig. 4f). This result is in agreement with the largely unaltered maps of NADs in senescence. In the next experiments the spatial dynamics of tandem repeat arrays was addressed, which build the core of the nucleolus-associated chromatin, but are not present on NAD maps because they reside in non-annotated genomic regions. The clusters of rDNA and the distal junctions of NORs displayed basically no difference between young and senescent cells. There was no indication for complete inactivation of NORs, that is, complete separation of strong hybridization signals from the nucleolus. The distal junctions appeared as discrete loci at the nucleolar periphery (Fig. 4g), whereas rDNA signals were distributed within the nucleoli (Fig. 4h). Similar to these loci, the global distribution of telomeres did not show detectable aging-related difference at the nucleolar and peri-nucleolar space, however the signal intensities were severely reduced in replicative senescent cells indicating telomere attrition (Fig. 4i). In contrast, senescence-associated distension of satellite repeats led to their substantial disassociation from nucleoli (Fig. 4j and 4k). Collectively, the 3D immuno-FISH and genomics analyses showed that mainly satellite repeat arrays and individual genes within NADs are the subjects of nucleolus-associated chromatin reorganization in senescence.

### Modification of H3K9 may represent a regulatory mechanism in senescence-dependent global chromatin alterations

In order to identify the epigenetic regulatory network underlying the senescence-dependent alterations of nuclear architecture, gene set enrichment analyses (GSEA) were performed (Fig. 5a and Supplementary Fig. S11). Comprehensive gene sets were compiled that include genes encoding ATPase and regulatory subunits of chromatin remodelling complexes (CRC), histone chaperones regulating chromatin assembly and disassembly, DNA modifying enzymes and proteins that bind modified DNA, furthermore histone modifying enzymes (Supplementary Table S5). The results revealed a global reduction of the mRNA levels of epigenetic regulators in senescence, which correlates well with the non-proliferative status of the cells. However, the transcript levels of a small number of genes, including *ERCC6, BAZ2A, KAT2B, SETDB2, EZH1, KDM3A* and *KDM7A,* were significantly increased. This strongly suggests that their gene products could be actively involved in genome regulation in replicative senescence. Notably, we also observed significant up- or down-regulation of various histone modifying enzymes that use H3K9 as substrate (Fig. 5a). The result suggests that this residue may act as target point of different senescence-dependent cellular signals.

**Figure 5.**
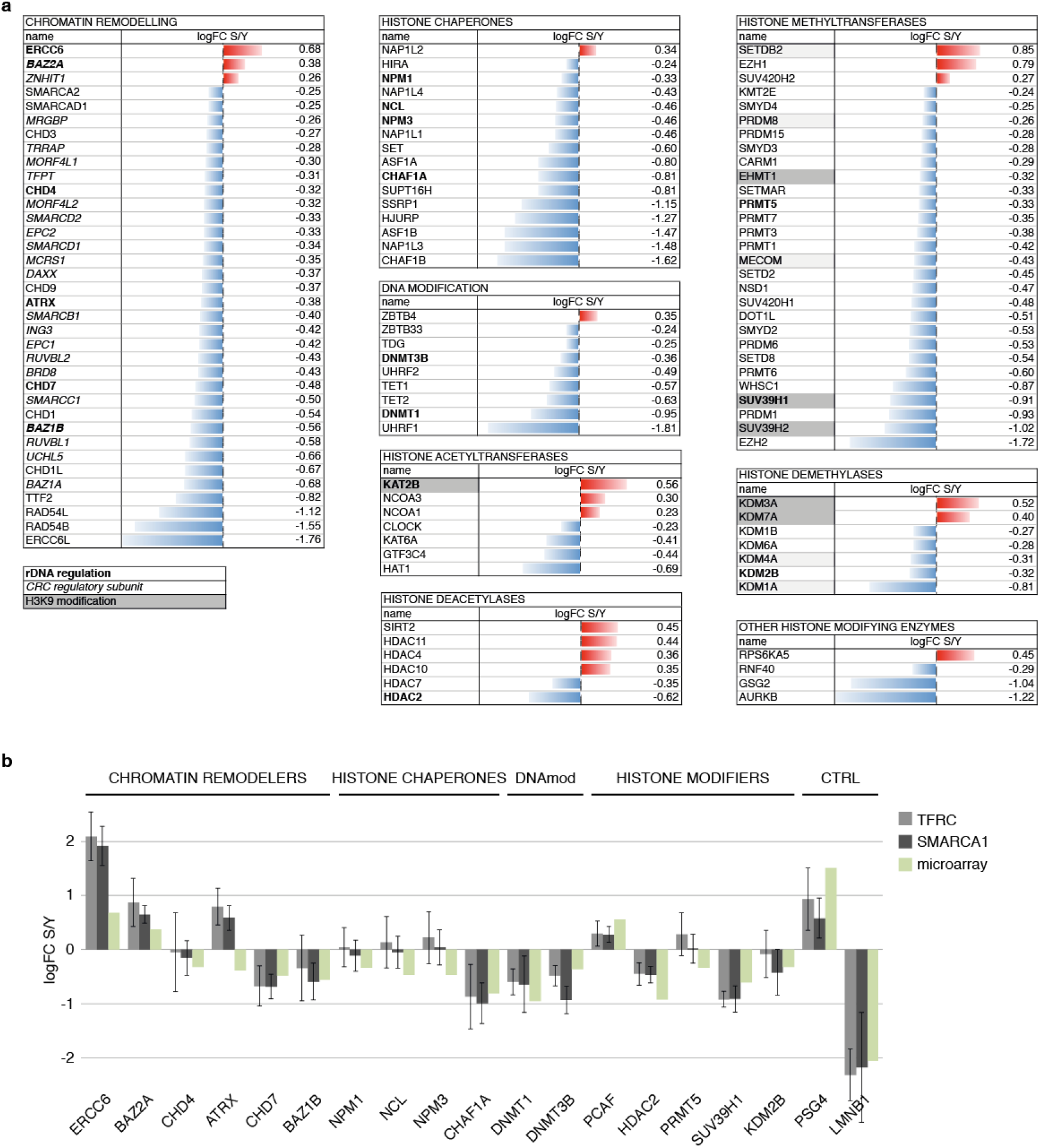
Gene set enrichment analysis (GSEA) identifies the epigenetic regulatory network of senescence-dependent chromatin alterations and suggests that the H3K9 residue is a control point in this process. (**a**) GSEA of epigenetic regulators (see Supplementary Table S5 for the full list) reveals frequent decrease (blue bars) in their mRNA levels in senescence. The few genes with increased mRNA levels, which are indicated in red, can be considered as active regulators of senescence. Bar graphs show log2 fold changes in mRNA levels of senescent vs. young cells. Epigenetic regulators with known nucleolar, rDNA-related activity are shown in bold, and regulatory subunits of chromatin remodelling complexes (CRC) in italic. Histone modifying enzymes that act certainly (n=6) or possibly (n=5) on H3K9 are labelled with dark and light grey background, respectively. (**b**) Quantitative RT-PCR validation of selected microarray data. Relative amounts of specific mRNA species in total RNA preparations from young and senescent cells were determined by quantitative RT-PCR with primer pairs listed in Supplementary Table S6. The bar graphs depict log2 fold changes in mRNA levels of senescent vs. young cells as determined using two different calibrator mRNA species in qRT-PCR experiments. The log2 fold changes measured in microarray experiments are shown next to the corresponding qRT-PCR data (see legend of the graph). Error bars represent the standard deviation of three independent biological replicate experiments, each of which was analysed in triplicate quantitative PCR reactions.

To further validate the results of gene expression array experiments, quantitative RT-PCR was performed (Fig. 5b and Supplementary Table S6). The levels of nineteen mRNA species, which showed significant alterations in the microarray analysis, were measured in proliferating and senescent cells, and two control mRNAs *(TFRC* and *SMARCA1)* were used for normalization. Seventeen targets *(ERCC6, BAZ2A, CHD4, ATRX, CHD7, BAZ1B, NPM1, NCL, NPM3, CHAF1A, DNMT1, DNMT3B, PCAF, HDAC2, PRMT5, SUV39H1, KDM2B)* were selected based on their known nucleolar function ^53^ The remaining two, *PSG4* and *LMNB1,* were selected because they show strong up- and down-regulation in senescence, respectively. The qRT-PCR results largely supported the results of the microarray analysis, as only the results for *ATRX* showed clear difference in the two independent assays.

### The nucleolus plays a key role in the senescence-dependent spatial dynamics of H3K9me3-marked chromatin organization

Next we decided to uncover senescence-dependent alterations of H3K9me3-marked constitutive heterochromatin in our model system for the following reasons: First, several modifiers of the H3K9 residue show significantly altered transcript levels in senescent compared to proliferating cells (Fig. 5). Second, the role of H3K9 methylation has been demonstrated in the organization of nucleolus-associated chromatin in *Drosophila* ^54^ Third, while genomic redistributions of H3K4me3 and H3K27me3 histone modifications, markers for active and facultative heterochromatin respectively, were investigated in replicative senescence in IMR90 cells ^26^, H3K9me3-marked chromatin dynamics was analysed only in oncogene-induced senescence ^24,55^ Fourth, H3K9me3-marked centromeric and pericentromeric satellite repeat clusters are essential components of nucleolus-associated chromatin 15, and they are subject to senescence-associated distension (see Swanson *et al.* ^29^ and Fig. 4j-k). Semi-quantitative immunoblot analyses were performed to monitor the overall level of H3K9me3, and the results showed that there is a strong decrease in senescent cells compared to young, proliferating ones. However, this change goes hand in hand with a somewhat less decrease in global histone H3 level. In addition, robust decrease in Lamin B1 and no detectable alteration in Lamin A/C levels were observed (Fig. 6a and Supplementary Fig. S12). Next, the co-localization of the strongest H3K9me3 signals with pericentromeric HSATII repeat clusters was demonstrated in 3D immuno-FISH experiments in IMR90 cells (Fig. 6b). In order to reveal senescence-associated changes in the nuclear distribution of constitutive heterochromatin, quantitative immunofluorescence experiments were performed. The analysis focused on the perinucleolar space and the nuclear periphery (Fig. 6c), the two main sites of heterochromatin accumulation in the human nucleus. First, the fluorescence intensity of the H3K9me3 staining was measured in the respective areas and calculated as per cent of the total fluorescence intensity. The results revealed that H3K9me3 signals decrease remarkably, from 5.2 % to 2.2 %, at the perinucleolar space in replicative senescence (Fig. 6d). In contrast, there was only a little difference between the relative fluorescence intensities of proliferating and senescent cells at the nuclear periphery (9.5 % vs. 8.4 %). Second, the coefficient of H3K9me3 signal variation (C.V. = standard deviation/mean of fluorescence intensity) was calculated, which gives an indication of the heterogeneity of the staining in the respective areas (Fig. 6e). The results showed a more heterogeneous staining in the nucleus of senescent IMR90 cells with 0.677 vs. 0.573 C.V. values in senescent and proliferating cells, respectively. Interestingly, at the perinucleolar space the C.V. values did not differ significantly in proliferating and senescent cells (0.576 compared to 0.555), but a significant increase was detected at the periphery in senescent compared to proliferating cells (0.709 vs. 0.632). Last, the abundance of the most heterochromatic regions in the perinucleolar space and at the nuclear periphery was evaluated. Therefore, the image containing the H3K9me3 signals was segmented by thresholding the fluorescence signals at 90% of the maximal intensity. While the abundance of the 10% brightest pixels, which represent the most heterochromatic regions, did not change significantly at the nuclear periphery (15.7% in proliferating vs. 13.7% in senescent cells), it strongly decreased at the perinucleolar space (5.6% in proliferating vs. 1.3% in senescent cells) (Fig. 6f). We suppose that the global loss of H3K9me3 at the perinucleolar space (Fig. 6d) is mainly due to the senescence-associated distension of satellites, which results also in the selective loss of intensely stained areas (Fig. 6f). In contrast, the signal heterogeneity remains largely unaltered at the perinucleolar space (Fig. 6e), which could be explained with the stable nucleolus association of NADs in senescence (Fig. 4 and Supplementary Fig. S9). Considering that the H3K9me3 signal distribution gets heterogeneous in other nuclear regions (Fig. 6e), we conclude that the nucleolus may play a functionally important role in maintaining the 3D organization of constitutive heterochromatin in replicative senescence.

**Figure 6.**
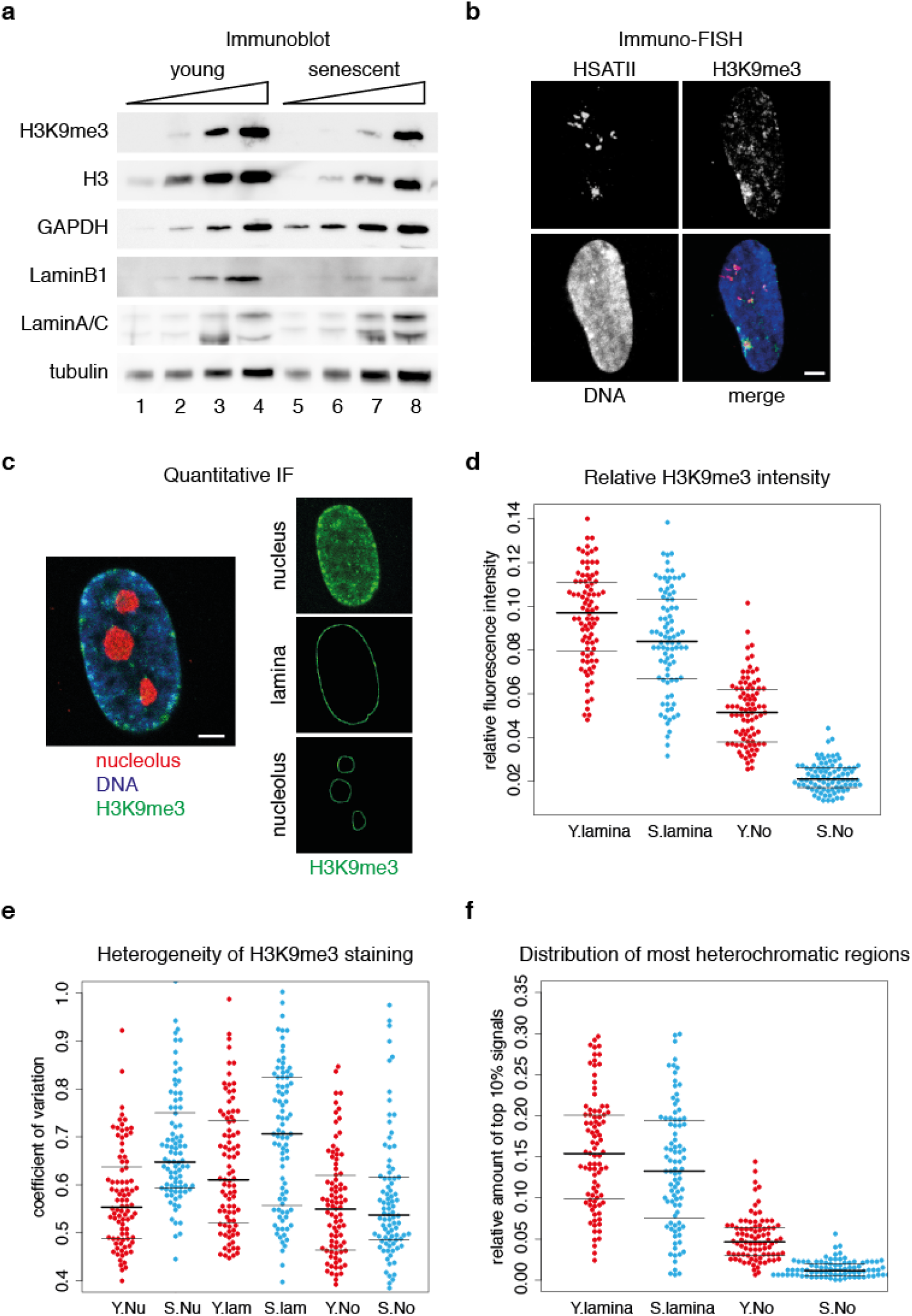
The nucleolus plays a key role in the senescence-dependent spatial dynamics of H3K9me3-marked chromatin organization. (**a**) Semi-quantitative immunoblots show severely decreased H3K9me3 and Lamin B1 levels, less strongly decreased H3 levels and no detectable alterations in Lamin A/C, tubulin and GAPDH levels in senescence. The same amounts of whole cell extracts of young and senescent cells were loaded as serial two-fold dilutions on SDS-PA gels and analysed on immunoblots (see the entire dataset from three independent experiments in Supplementary Fig. S10). (**b**) H3K9me3 is accumulated at spatially compact satellite repeat clusters. 3D immuno-FISH shows strong co-localization of H3K9me3 and HSATII staining. Mid-section of a representative confocal microscopy image is shown. HSATII FISH signals are in red, DAPI counterstain in blue, and H3K9me3 immunofluorescence signals are in green (scale bar: 1.6 μm). (**c**) Quantitative immunofluorescence analysis of H3K9me3 distribution. The areas of interest are illustrated on a light optical section of a representative confocal microscopy image. The lamina- and nucleolus-associated areas label 240 nm distances from the edges of the DAPI and nucleolus staining, respectively. (**d**) Bee swarm plots of relative fluorescence intensities show senescence-dependent small decrease in H3K9me3 levels at the nuclear periphery and strong reduction at the perinucleolar space. Proliferating and senescent IMR90 cells were stained for H3K9me3 and the relative immunofluorescence intensities were measured at the nuclear periphery (lamina) and at the perinucleolar space (No). Values measured in proliferating cells ('Y') are shown in red, values measured in senescent cells ('S') are shown in blue. Results from individual cells are illustrated as single data points (n_Y_=88, n_S_=88). A solid line indicates the median, and thin lines the upper and lower quartile. Median: Y.lamina = 0.095, S.lamina = 0.084; Y.No = 0.052, S.No = 0.022. (**e**) Bee swarm plots indicate more heterogeneous H3K9me3 staining in the nucleus and at the nuclear periphery of senescent cells, but no change in the perinucleolar space. The heterogeneity of staining was calculated as coefficient of variation (C.V. = standard deviation/mean of fluorescence intensity) for the total nucleus (Nu), the nuclear periphery (lamina) and the perinucleolar space (No). Plot labels are as in (**d**). Median: Y.Nu = 0.573, S.Nu = 0.677; Y.lamina = 0.632, S.lamina = 0.709; Y.No = 0.576, S.No = 0.555, n_Y_=88, n_S_=88. (**f**) Bee swarm plots illustrate robust rearrangement of the most heterochromatic regions in the perinucleolar space. The distribution of the 10% brightest pixels was quantified at the nuclear periphery (lamina) and the perinucleolar space (No). Ratios were calculated compared to the whole nucleus. Plot labels are as in (**d**). Median: Y.lamina = 0.157, S.lamina = 0.137; Y.No = 0.056, S.No = 0.013. n_Y_=88, n_S_=88.

## Discussion

In this study we used cell fractionation and genomics analyses to discover the features of NADs in proliferating and replicative senescent human embryonic fibroblasts and to untangle their network. Our results demonstrate that NADs are predominantly heterochromatic chromosomal domains, which remain largely associated with nucleoli also in senescent cells. We show that, despite microscopically observable nuclear and nucleolar re-organization, only sub-NADs change their association with nucleoli during cellular aging. These changes clearly correlate with altered transcriptional activity. Our data suggests that transcriptional activity-dependent disassociation is characteristic also for X escaper genes ^43^ located on the nucleolus-associated inactive X chromosome ^56,57^ The largest senescence-dependent rearrangements of the nucleolus-associated chromatin are, however, not visible on the genomic maps of NADs, because they are caused by the distension of satellite repeat clusters that are not present in genome assemblies. We show altered nucleolar association of pericentromeric and centromeric satellite repeats, which corresponds to the SADS phenotype. Notably, SADS was shown to coincide with increased transcription of satellite repeats ^58^ T o summarize these findings we propose a model in which active NORs, bearing active and potentially also inactive rDNA repeats ^59^ and additional NOR-specific sequence elements around the rDNA clusters ^60^, safeguard the maintenance of the NAD network in replicative senescence. At the same time, transcriptionally activated regions of specific NADs and satellite repeat clusters get disassociated from nucleoli and consequently alter the composition of nucleolus-associated chromatin. We screened for potential epigenetic regulators of this process by using a GSEA approach and observed the high prevalence of H3K9 modifying enzymes on the list of genes with significantly altered expression level. Concordantly, we demonstrate global loss and local spatial rearrangements of H3K9me3-marked heterochromatin in replicative senescence. These results not only support each other, but also reveal a role for this histone modification in the senescence-dependent dynamics of nucleolus-associated chromatin. Remarkably, we found also *BAZ2A* among the handful of chromatin regulator genes that show elevated expression level in senescence. The BAZ2A protein is thought to be involved in silencing and constitutive heterochromatin formation at rDNA, centromeres and telomeres ^61,62^, and it has been recently reported that depletion of BAZ2A promotes escape from senescence ^63^. This suggests a role for BAZ2A in the regulation of a subset of H3K9me3-marked chromatin domains in senescence.

Although senescence-dependent chromatin alterations have been investigated in recent studies ^24^•^28,55^, several aspects of the nuclear changes remained unclear. One of them, the dynamics of nucleolus-associated chromatin, has been addressed here. Still, the picture is far from complete, and differences in the experimental systems make direct comparisons between the different studies often difficult. For instance, SAHF formation was not prominent in our replicative senescence experimental setup, which distinguishes this study from comprehensive chromatin and nuclear architecture analyses of oncogene-induced senescence (OIS) with a characteristic SAHF phenotype ^24,27,55^. Nevertheless, the changes observed in constitutive heterochromatin rearrangements in replicative senescence in our work are partially consistent with the results of OIS and other replicative senescence studies. Most importantly, the alterations of H3K9me3 patterns at the nuclear periphery are reminiscent of its previously reported reorganization in LADs ^24^ This reorganization accompanies senescent-dependent Lamin B1 depletion, which was demonstrated also here. Notably, we find about 70% overlap between the genomic maps of NADs and LADs in human fibroblasts, which is in agreement with the stochastic association of several chromosomal domains with either the lamina or the nucleolus ^64,65^, and in general with the probabilistic model of the spatial organization of the human genome ^3^ Indeed, when nucleoli are in close proximity to the lamina, several chromosomal domains may also be simultaneously associated with both nuclear compartments. We predict that NADs on chromosomes bearing an active NOR show higher contact frequencies with the nucleolus than with the lamina. Our chromosome-based NAD-LAD comparison, and a pioneering single-cell genomic analysis of LADs in KBM7 cells seem to support this idea ^66^ However, this issue should be addressed by single-cell nucleolus genomics in future studies, in which the dissimilar senescence-dependent dynamics of LADs and NADs could also be more precisely addressed. We hypothesize that globally lowered contact frequencies with the nuclear periphery may accompany with increased nucleolar contact frequencies. This scenario would be consistent with the senescence-dependent destruction of the lamina and the simultaneous enlargement of nucleoli. Notably, the latter one correlates with increased ribosomal RNA precursor levels, which is due to delayed processing and acts as a senescence inducer ^67^ In a next step, in nuclei undergoing SAHF formation the association probability of chromosomal domains with the nucleolus might also be reduced.

Concerning the senescence-independent role of NADs in chromosome organization, bioinformatics analyses of high-resolution IMR90 Hi-C data ^48^ reveal that NAD-NAD interactions are the dominant ones in the 10-50 Mb genomic distance range and in intra-chromosome-arm contacts, whereas iNAD-iNAD interactions are more frequent over larger genomic distances. The high interaction frequency of NADs in the 10-50 Mb distance range correlates well with the current view that a compact, transcriptionally inactive nuclear compartment builds the core of chromatin domain clusters ^68^ Importantly, the pattern of NADs and iNADs largely resembles that of the TADs, which connects the spatial organization of the interphase genome to the nucleolus, the centre of ribosome biogenesis. As ribosomal RNA synthesis determines nucleolar assembly and represents the primary response site of cell growth regulation, we speculate that active NORs may physically link the cell's metabolic activity to 3D genome organization. Notably, the spatial organization of the genome can be considered as moderator of chromosomal communication ^69^ The presented study and several previous observations about the role of the nucleolus in shaping genome architecture from yeast to human (reviewed in ^12^^14^,^70^,^71^) lead us to postulate that the nucleolus is an important factor in moderating chromosomal communication.

## Methods

### Cell culture and nucleolus isolation

Human IMR90 embryonic fibroblasts were obtained from Coriell Repositories (Cat. No. I90-79) and cultivated in DMEM (Gibco Cat. No. 21885-025 supplemented with 10% v/v Foetal Calf Serum, 100 U/mL Penicillin, 100 μg/mL Streptomycin) at 37°C in humidified, 5% CO_2_ atmosphere and regularly tested for mycoplasma contamination. Proliferating young and senescent cells were cross-linked with 1% formaldehyde and nucleoli were isolated as described^72^. Nucleolus-associated DNA was prepared from two independent experiments, both from young and senescent cells, for subsequent microarray analysis. Total genomic DNA of the same IMR90 cells served as control for one young and one senescent sample during comparative hybridization, whereas the other young and senescent samples were hybridized against non-nucleolar DNA. This DNA was collected during the nucleolus preparation and only the DNA of the last, nucleolar fraction was excluded from it. Since the depletion of nucleolus-associated DNA in the control did not alter markedly the NAD patterns, the results obtained from the two independent experiments were combined both for the young and senescent samples (Supplementary Fig. S1). Quality controls of nucleolus preparations were performed as described^15^.

### Cellular senescence assays

Cell populations were kept in culture for two weeks after they stopped growing. During this time the growth medium was replaced every second day. The senescence status of the IMR90 cell populations was monitored by senescence-associated beta-galactosidase staining of fixed cells (BioVision Senescence Detecion Kit), and immunofluorescence staining of the MKI67 proliferation marker protein in fixed cells by using a rabbit polyclonal antibody (Santa Cruz sc-15402).

### Computational analyses

If not stated otherwise, all computational analyses were performed in R/bioconductor using default parameters (R-project.org/bioconductor.org, version 3.3; R Core Team (2016). R: A language and environment for statistical computing. R Foundation for Statistical Computing, Vienna, Austria. URL https://www.R-project.org/). The hg19 version of the human genome served as reference. We used hg19, because most of the comparative genomics analyses were performed with datasets available in this format.

### Gene expression microarray experiments

Gene expression microarray analyses were performed using TRIzol-extracted total RNA from young, proliferating and senescent IMR90 human diploid fibroblasts. Two biological replicate experiments were performed, and the Affymetrix Human Gene 1.0 ST microarray platform was used. Labelling of the samples and hybridizations were carried out at Source BioScience. We calculated expression values using the Robust Multichip Average (RMA) algorithm. Many-to-one probesets to gene relationships were resolved by retaining the probeset with the highest variance across all arrays. In order to compare expression levels with the ones of Shah et al. (GSE36640) and Lackner et al. (E-MTAB-2086) we merged all experiments using the 'COMBAT' method in the bioconductor library 'inSilicoMerging'. Hierarchical clustering on the merged set was performed on euclidean distances with the 'complete' method of the 'hclust' function in R.

### Gene Set Enrichment Analysis

Gene sets of chromatin remodelling enzymes, histone chaperones, DNA modification enzymes and binding proteins, furthermore histone modifying enzymes were compiled based on literature search. Gene Set Enrichment Analyses (GSEA) were performed according to the instructions described on the GSEA homepage http://software.broadinstitute.org/gsea/index.jsp and in ^73^

### Quantitative RT-PCR

To validate the results of the gene expression microarray experiments, RNA of young and senescent cells was prepared from three additional, independent experiments. 200 ng RNA of each sample was reverse transcribed by using 200U of MMLV-RT and random hexamers. The resulting cDNA was amplified in real-time quantitative PCR experiments using SybrGreen-intercalation-based quantification. Primer pairs used for amplification were selected using the qPrimerDepot (http://primerdepot.nci.nih.gov). Data were collected with a Rotor-Gene Q system (Qiagen) and analysed using the comparative quantitation module of the system software. The mean and standard deviation values are derived from three independent experiments analysed in triplicate quantitative PCR reactions.

### Mapping and genomics of NADs

Raw tiling array signals were subjected to quantile normalization. Nucleolar enrichment was defined as the log2-fold difference of the nucleolar signal over the background (genomic input or supernatant, respectively). Enrichment signals were smoothed per sample by sliding medians in 100 kb windows as in our previous study ^15^ Subsequently, enrichments were averaged for proliferating and senescent cells, respectively. Given the bimodal nature of signal distribution, a two-state hidden Markov model (library 'tileHMM') was employed to classify nucleolus-associated domains with a minimum length of 10 kb (Supplementary Fig. S2). The average enrichment values within each domain served as 'NAD-scores'. Genomic features of NADs and iNADs were determined by using the UCSC Table Browser (http://genome.ucsc.edu/cgi-bin/hgTables). Jaccard similarity indices that reflect the ratio of the number of intersecting base pairs between two sets to the number of base pairs in the union of the two sets and thus serve as a similarity statistic for comparing the distribution of regions of different genomic features were calculated with bedtools' 'jaccard' function (version 2.24).

### Hi-C data analysis

Raw inter- and intracromosomal contact matrices (10 kb windows, MAPQ>=30) for the IMR90 map presented in ^48^ (GSE63525) were normalized using the provided normalization vectors. Each interaction was classified for interaction distance, overlap with NAD or iNAD, and chromosomal arm localization.

### 3D immuno-FISH

The 3D FISH experiments, and subsequent confocal microscopy and image analysis were essentially performed as described ^15^, except that series of optical sections through 3D-preserved nuclei were collected using a Leica TCS SP8 confocal system. In the localisation experiments anti-NPM1 (Santa Cruz sc-6013R or sc-56622) and different fluorescence dye-conjugated secondary antibodies, BAC clones RP11-81M8, RP11-123G19, RP11-89O2, RP11-828F4, RP11-125O21, RP5-915N17, RP11-89H10, RP11-44B13, RP11-434B14, RP11-413F20, RP11-1137G4, RP11-173M10, the 'DJ' cosmid clone LA14 138F10 ^60^, the pHr4 plasmid DNA containing the +18063/+30486 BamHI/EcoRI intergenic spacer fragment of the human rDNA (GenBank Acc. No. U13369) in pBluescript SK+ ^74^, furthermore 5'-biotin labelled LNA FISH probes for human HSATII, telomere and centromeric alpha-satellite detection (Exiqon) were used.

### Immunofluorescence

Cells were rinsed twice in 1xPBS and fixed in 4% paraformaldehyde in 1xPBS for 10 min. All steps were carried out at room temperature. During the last minute of fixation, few drops of 0.5% TritonX-100 in PBS were added. Cells were then washed three times in 1xPBS/0.01% Triton X-100 for 3 min, followed by 5 min in 1xPBS/0.5% Triton X-100. Finally, cells were washed twice with PBS-T (1xPBS/0.1% Tween20) for each 5 min before antibody staining. Cells were incubated with the primary antibodies (anti-NPM1: Santa Cruz sc-6013R, anti-H3K9me3: CMA318^75^, anti-H3K27me3: Active Motif 61017, anti-LMNB1: Santa Cruz sc-6216) in 4% bovine serum albumin (BSA) in PBS-T for 1h in a humidified chamber, washed in PBS-T three times for 3 min and incubated with fluorescence dye-conjugated secondary antibodies in 4% BSA in PBS-T for 1 h. After washing with PBS-T twice for 5 min, the DNA was counterstained with 50 ng/ml of DAPI in PBS-T for 5 min. Slides were rinsed in PBS-T and mounted in Vectashield (Vector). Middle optical sections of 200 nm z-stacks were selected from 8-bit CLSM (Leica TCS SP8) images based on the quality of nuclear morphology. The xy pixel size was 80.25x80.25 nm. A custom-made ImageJ script was used for semi-automated quantitative analysis of the immunofluorescence signals^76^. The following parameters were measured: i) The percentage of total nuclear fluorescence intensities of H3K9me3 within the lamina- and nucleolus-associated areas (regions of interests - ROIs), which label 240 nm distance from the edge of the DAPI and nucleolus staining, respectively, ii) The heterogeneity of the staining in the ROIs as coefficient of variation (C.V. = standard deviation/mean of fluorescence intensity^77^), iii) The percentage of the 10% brightest nuclear pixels in the two ROIs. Nucleolar number and volume measurements were performed on z-stack images acquired with a Zeiss Axiovert200 microscope, using a 'Plan-Apochromat 63x/1.40 Oil' objective. Whole nuclei were recorded with a z-step size of 500 nm. Nucleoli were counted and assigned to cells manually. Voxel sizes were obtained by calibrating the objective, and values were saved and transferred to FIJI/ImageJ and the volume in μm^3^ was calculated by measuring the number of object voxels using the '3D Objects Counter' plug-in. The measurements were performed for both the DAPI and nucleolus staining to determine nuclear and nucleolar volumes, respectively. The resulting values were transferred to an Excel sheet and the average nucleolar volume, the total nucleolar volume per cell and the ratio of nucleolar/nuclear volume per cell wascalculated. The results were displayed as bee swarm plots, on which the data points represent individual cells.

### Western blot

Whole cell extracts from the same amounts of young and proliferating cells were prepared, separated by SDS-PAGE and blotted by semidry transfer. The western blot membranes were incubated with antibodies against GAPDH (Cell Signaling, #5174), histone H3 (Abcam, ab1791), the histone modification H3K9me3 (CMA318, ^75^), tubulin (Abcam ab7291), Lamin A/C (Santa Cruz, sc-20681) and Lamin B1 (Santa Cruz, sc-6216).

### Data availability

Array CGH data used for NAD mapping and gene expression microarray data have been submitted to the NCBI Gene Expression Omnibus (GEO; http://www.ncbi.nlm.nih.gov/geo/) under accession numbers GSE78043 and GSE80447, respectively.

## Acknowledgements

We thank Gernot Längst, Herbert Tschochner, Philipp Milkereit and Joachim Griesenbeck for infrastructural and financial support, Thomas Cremer for critical reading of the manuscript, Thomas Dresselhaus for confocal microscopy access, Hiroshi Kimura for providing H3K9me3 antibodies, Brian McStay for providing the LA14 138F10 cosmid and pHr4 plasmid, furthermore Regina Gröbner-Ferreira, Sebastian Fladerer, Christina Fischer and Florian Hirsch for technical help. This work was supported by the DFG SFB960 programme.

## Author contributions

SD and AN performed all wet-lab experiments, furthermore analysed and interpreted the data of single-cell experiments. TS conducted all computational biology analyses with the exception of the GSEA, which was performed by AN. AN designed the study, TS contributed to the design of bioinformatics analyses. AN wrote, TS and SD commented the manuscript. All authors read and approved the final manuscript.

